# Key residues in SARS-CoV-2 NSP3 hyper variable region are necessary to modulate early stress granule activity

**DOI:** 10.1101/2025.11.22.689758

**Authors:** R. Elias Alvarado, Kumari G. Lokugamage, Dimitriya Garvanska, Leah K. Estes, Yani Ahearn, Alyssa M. McLeland, Arian Moayyed, Jennifer Chen, Blanca Lopez Mendez, Jessica A. Plante, Kenneth S. Plante, Bryan A. Johnson, Jakob Nilsson, Vineet D. Menachery

## Abstract

Antagonism of the host responses that limits viral replication is critical to the success of infection. Recently, we identified that the hypervariable region (HVR) of SARS-CoV-2 NSP3 binds to FXR1 and disrupts stress granule formation during the early stages of infection. Despite variation across the rest of the HVR, a 20-amino acid region, highly conserved in the Sarbecovirus family, is required for NSP3-FXR1 binding, but the critical residues remained unresolved. In this study, we explore the individual residues in NSP3 driving FXR1 binding and determine their impact on viral replication, pathogenesis, and stress granule formation. Our results indicate that the tyrosine at position 138 (Y138) and a phenylalanine at position 145 (F145) are required for FXR1 binding and affinity. Using reverse genetics, we showed mutating NSP3 Y138A/F145A (YF mutant) reduced viral replication *in vitro* and *in vivo.* Importantly, we demonstrate that attenuation is not due to differential type I interferon responses, but rather loss of stress granule control by the NSP3 mutant as compared to WT. Together, our findings demonstrate the importance of Y138 and F145 within the NSP3-HVR in regulating stress granule formation at the early times post infection.

**IMPORTANCE:** Stress granules play a key role in host-antiviral defenses and viruses have developed strategies to antagonize their activity. For SARS-CoV-2, the virus has two proteins that antagonize stress granules with NSP3 acting early and nucleocapsid acting at late times. Here, we show that key NSP3 residues Y138 and F145, conserved across the Sarbecovirus family, are necessary to bind FXR1 and disrupt its activity in stress granule formation. Mutating these residues results in attenuation of SARS-CoV-2 replication and induces stress granule formation at early times post infection. These results show the importance of these NSP3 residues in disrupting stress granule formation early and highlight multiple approaches SARS-CoV-2 uses to antagonize stress granule activation.

## INTRODUCTION

SARS-CoV-2 is the causative agent of COVID-19 and is responsible for an unprecedented six million deaths worldwide^1^. Despite our incredible strides in medicine and research, our understanding of how SARS-CoV-2 induces disease and evades immunity remains limited. To solve this problem, future research will require a molecular understanding of events driving pathogenesis and viral replication.

SARS-CoV-2 is a positive-sense, single-stranded RNA viruses with a genome size approximately 30kb ^2^ The viral genome contains two open reading frames (ORFs) that encode two polyproteins; polyprotein 1A and 1AB. After translation, the polyproteins are cleaved into their individual nonstructural proteins (NSPs) components ^3^. Overall, the SARS-CoV-2 genome encodes 16 nonstructural proteins (NSP 1-16), 4 structural and 9 accessory proteins. Extensive research has shown that NSPs play critical roles in supporting viral replication while simultaneously evading innate immune responses. Amongst them, NSP3 is the largest protein that contains several conserved and non-conserved domains that are exclusive to *Sarbecovirus* ^4,5^. At the N-terminus of NSP3 lies a unique region rich in Glu/Asp-amino acids termed the hypervariable region (HVR)^6^. Despite being a region that exists in all CoVs, the function of the HVR has yet to be determined. Being an intrinsically disordered region (IDR), the HVR harbors short (3-10 residue) fixed amino acid sequences called short linear motifs (SLiMs) which can serve as important sites for protein-protein interactions.^7^ Therefore, the HVR has the potential to serve as a hub for binding to host cellular factors during infection.

Foreign stressors, including virus infections, trigger cells to temporarily halt protein translation to focus efforts on survival and return to homeostatic conditions. Stress Granule (SG) formation is one response that occurs in the context of host translation arrest and is frequently observed during viral infections ^8^. SGs, membrane-less cytoplasmic structures, are large focal complexes composed of untranslated mRNAs, protein translation factors, and RNA-binding proteins (RBPs) ^9,10^. Upon activation, SG components like G3BP1, FXR proteins, and TIA-1 form condensates through liquid-liquid phase separation that is driven by multivalent interactions between the granule constituents^9,11^. When these liquid droplets form, untranslated mRNA and protein translation factors are sequestered, impairing translation. Once the stressor is gone, SGs disassemble, and release these factors allowing translation to proceed. Importantly, viral replication is dependent on the host translational machinery and thus SGs play a role in host antiviral responses. Therefore, viruses have evolved ways to inhibit SGs by targeting key SG components, thereby disrupting the formation or facilitating disassembly of SGs during infection.

Among the key SG components, G3BP1 is an RNA binding protein that is the central regulator for SG formation^12,13^. G3BP1 plays a critical role in the dynamics of SGs via its contributions to liquid-liquid phase separation^14^. Importantly, G3BP1 is a common target of viral antagonism which often serves to disrupt SG activity following infection^15–19^. Our group and others have identified SARS-CoV-2 nucleocapsid as binding to G3BP1 and disrupting SG activity following infection ^20^. However, other RBPs including the FXR family of proteins have been implicated in SG formation and can be targeted by viruses during infection^21–24^. Associated with Fragile X Syndrome and other genetic diseases^25^, FXR proteins have a wide range of regulatory functions including formation of SGs. Utilizing the intrinsically disordered K-homology 2 (KH2) domain, FXR proteins can undergo liquid-liquid phase separation, which likely contributes to their role in SG formation ^26,27^. Importantly, viruses including old world (EEEV) and New world (VEEV) alphaviruses, adenovirus, and recently coronaviruses have also been shown to antagonize FXR function during infection ^23,28,29^. Given the reports of antagonism across multiple virus families, these results argue that there is a critical role for FXR proteins in SG activities and antiviral function.

Recently, our group identified that the HVR in NSP3 of SARS-CoV-2 acts as a hub to bind and hijack FXR1 away from SGs during early stages of infection ^30^. We show that cells expressing NSP3-HVR hijacks FXR1 away from SG but do not affect the overall number of SGs formed. Despite these findings in protein expressing cells, we cannot exclude that FXR1 and NSP3 interactions are important during actual viral infection. To further understand if the HVR of NSP3 is important for the regulation of SGs, we sought to refine our NSP3 viral mutant and further define the role for FXR1 during infection. Here, we found that tyrosine-138 (Y138) and phenylalanine-145 (F145) are key residues that are important for FXR1 binding and affinity. Using our reverse genetics system, we found SARS-CoV-2 mutants incapable of binding to FXR1 (YF mutant) show reduced viral replication *in vitro*, reduced viral load *in vivo*, and that this attenuation is not driven by type-1 interferon responses. Mechanistically, we show that preventing FXR1 binding to the HVR is important for recovering SG formation and that FXR1 acts as a proviral factor to support efficient viral replication. Our findings reveal that the NSP3-HVR is important for regulating SG formation at the early phase of infection to facilitate viral replication.

## Results

### NSP3 F145 and Y138 key to binding to FXR1

Our prior studies showed that SARS-CoV-2 NSP3 amino acid region 129-149 interacts with human FXR1 and that two SARS-CoV-2 alanine scanning mutants covering the 20 AA region were attenuated ^30^. The alanine scanning results revealed that multiple residues are responsible for the interaction. Additionally, a subsequent peptide array suggested a critical role for Y138 and F145 ^30^. Examining the AlphaFold based structure (**Fig. 1A**), the binding interface between FXR1 and NSP3 shows an interaction between FXR1 I304 and NSP3 F145 (**Fig. 1B**). While Y138 is not directly involved in the binding interface, the aromatic residue may offer stability to the loop allowing the FXR1/NSP3 interaction. To verify the role of these amino acids in binding, we generated a WT and Y138A/F145A mutant YFP-NSP3 fusion peptide (NSP3 1-181AA) (**Fig. 1C**). The YFP-NSP3 fusion proteins were transfected, lysates cleared and incubated with GFP-trap beads; following wash, samples were processed by western blot for FXR1 and GFP (**Fig. 1D**). As expected, GFP-trap lysates resulted in the pull down of FXR1 with WT YFP-NSP3 peptide; however, mutations Y138A/F145A ablated NSP3 binding to FXR1 demonstrating the importance of both residues for the FXR1/NSP3 interaction. We subsequently used isothermal titration to examine the molecular interactions between WT and mutant NSP3 peptide with FXR1 (**Fig. 2A)**. Injecting WT YFP-NSP3 peptide, we observed robust binding (2.2uM) to FXR1 (**Fig. 2B);** these results were consistent with our previous results^30^. In contrast, the mutation at Y138A/F145A in the NSP3 mutant peptide ablated binding to FXR1 (**Fig. 2C**). Together, these findings demonstrate that both Y138 and F145 in SARS-CoV-2 NSP3 are necessary for the FXR1 binding and their absence ablates the interaction.

**Figure 1:**
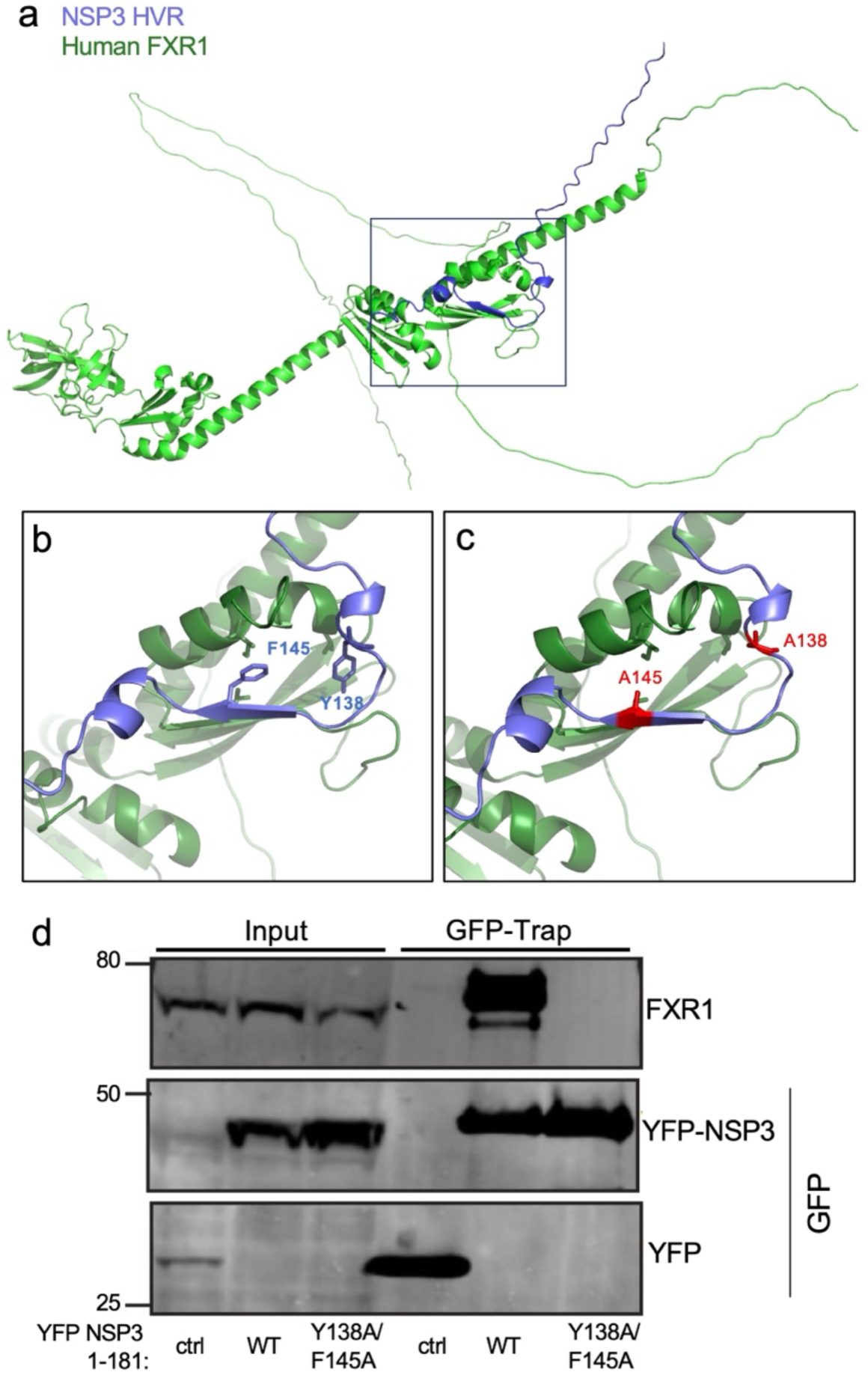
NSP3 mutations at Y138 and F145 disrupts peptide interactions with FXR1. (A) AlphaFold model of the FXR1-NSP3 complex interactions. (B) Purple indicated wild type NSP3 aromatic residues (C) while red indicate residues changed to alanine. (D)The indicated NSP3 fragments were fused to YFP, expressed, and immunoprecipitated from HeLa cells to examine binding to FXR1 by immunoblotting. Represented are two biological replicates.

**Figure 2:**
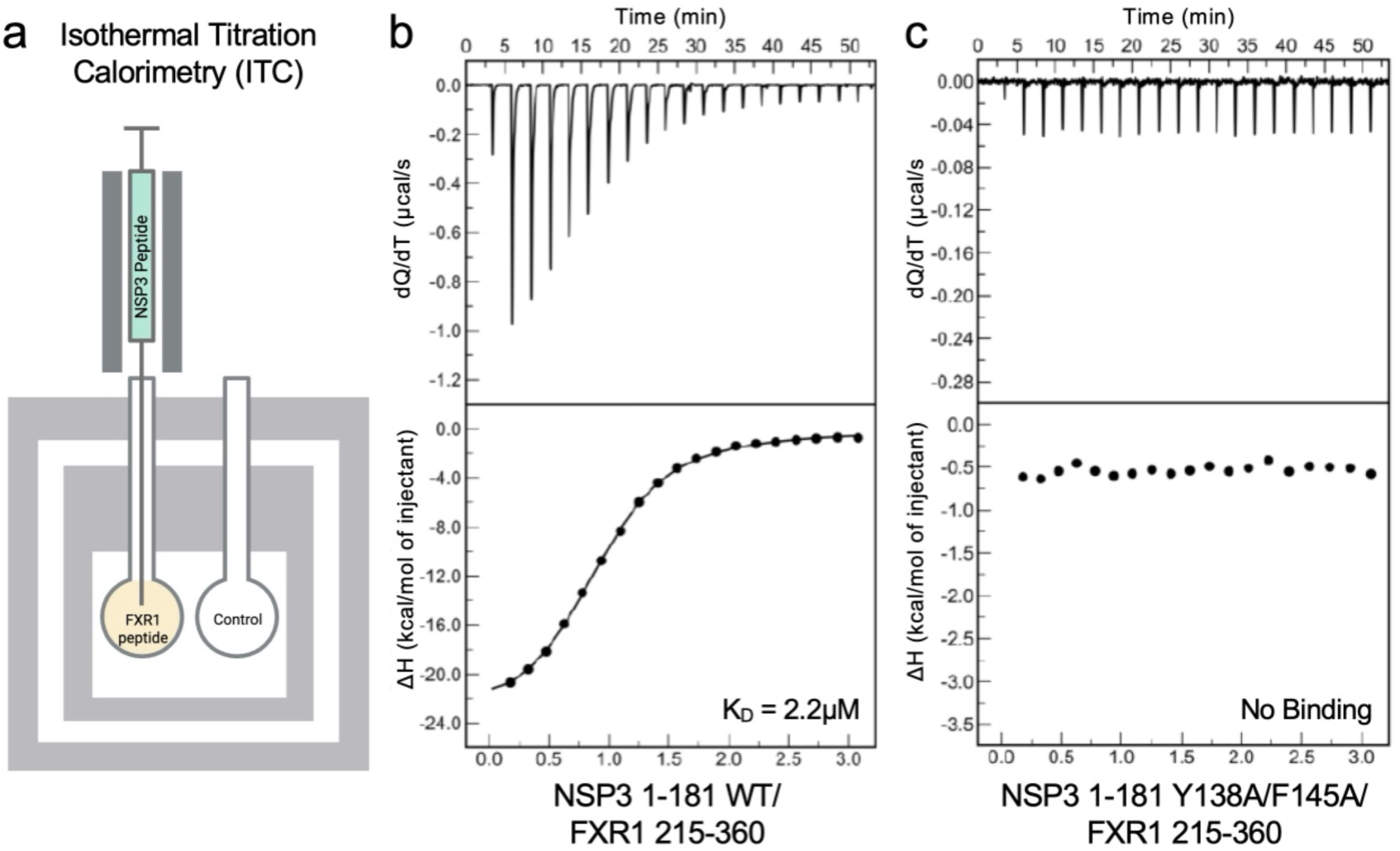
YF mutations ablate binding between NSP3 and FXR1. (A) Schematic of isothermal titration with injection of WT or mutant NSP3 peptide into chamber with human FXR1 peptide. A calorimeter measures the change in heat as the NSP3 peptide is titrated to calculate binding affinity (K_D_). Figure made using Biorender. B-C) Isothermal titration results between WT NSP3 peptide (B) and NSP3-YF mutant peptide (C) shows ablation of binding in the YF mutant compared to WT NSP3.

### Disrupting FXR1 binding to NSP3-HVR reduces viral replication *in vitro*

Y138 and F145 are both harbored within the hypervariable region (HVR) of NSP3, the largest viral protein produced during infection (**Fig. 3A**). To determine if these residues are conserved, we conducted a sequence alignment of NSP3-HVR from across *Sarbecovirus* members (**Fig. 3B**). Both Y138 and F145 were conserved in all *Sarbecoviruses* surveyed; however, similar residues or motifs could not be aligned from the NSP3 HVR of CoVs from the other alpha and beta CoV families. Interestingly, bat derived CoVs BANAL-236 and RATG3, the most SARS-CoV-2 similar sequences, retain all 20 residues in the FXR1 binding motif. In contrast, the bat CoVs RsSHC014 and WIV1, more closely related to SARS-CoV, have 6-7 residue differences compared to SARS-CoV-2 consensus. Overall, despite variation in the NSP3 HVR, Y138 and F145 are completely conserved across the entire Sarbecovirus family.

**Figure 3:**
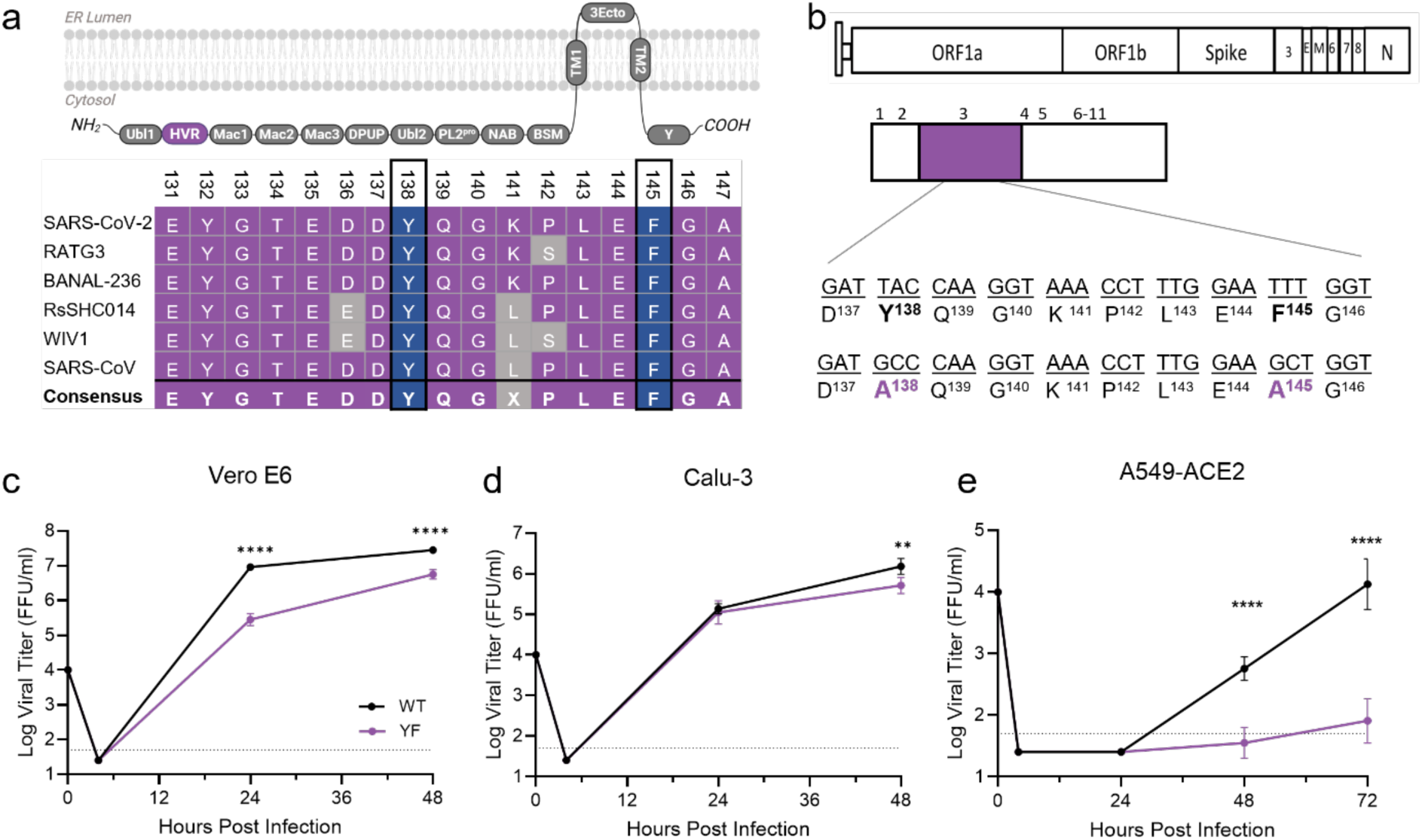
SARS-CoV-2 YF mutant has attenuated viral replication *in vitro*. (A) Schematic of the NSP3 domains including the hyper variable region (HVR, purple) made using Biorender. Sequence alignment of *Sarbecovirus* members NSP3-HVR shows conservation of Y138 and F145 across virus family. (B) Schematic of SARS-CoV-2 NSP3 with YF mutations to alanine (purple) introduced in Washington 1(WA-1) strain (C-E) Viral titer from Vero E6 (D), Calu3 (E), or A549-ACE2 (F) cells infected with WT (black) or YF (purple) SARS-CoV-2 at an MOI of 0.01 (n = 6 from two experiments, each with three biological replicates). Data are representative of mean ± SD. Statistical analysis measured by two-tailed Student’s t test. *P ≤ 0.05; **P ≤ 0.01; ***P ≤ 0.001, ****p<0.0001.

Having demonstrated that residues Y138 and F145 are necessary for binding with FXR1 (**Fig. 1** & **2**), we next evaluated the effects of disrupting this interaction during authentic viral infection. Here, we generated a SARS-CoV-2 NSP3 mutant that substitutes Y138 and F145 to alanine (YF) into the WT WA-1 backbone using our reverse genetics system ^31,32^ (**Fig. 3C**). Introducing the YF mutations caused no deficits in viral stock titer or changes in plaque morphology. We next compared the viral replication kinetics of YF mutant to WT across different cell lines using a low multiplicity of infection (MOI). We started with infection of VeroE6 cells, an African Green Monkey kidney cell line that lacks competent Type I interferon (IFN) responses (**Fig. 3D**). Following infection, the YF mutant had significant attenuation at both 24- and 48-hours post infection (HPI). We next evaluated the YF mutant in Calu3-2B4 cells, a human bronchial epithelial cell line with robust type I interferon responses ^33^. Following low MOI (0.01) infection, the YF mutant replicated to similar viral titers as WT at 24 HPI (**Fig. 3E**). By 48 hours, the YF mutant had a significant, but modest reduction in viral yields as compared to WT control. These results were contrast to VeroE6 results and demonstrated that attenuation of the YF mutant varied across different cell lines *in vitro*.

While we often observe a discrepancy between VeroE6 and Calu3 cells in regard to SARS-CoV-2 mutant replication, these differences often correlate with either the expression of TMPRSS2 or type I IFN response, both absent in Vero cells ^34,35^. With the opposite pattern, it argued attenuation may be driven by different factors including stress granule activity or control. To further examine the role of SGs, we examined A549-hACE2 cells, a lung adenocarcinoma cell line that are transduced to stably express human ACE2 receptor. Prior studies found robust SG responses are induced in A549 cells following virus infection and numerous SG assays had been evaluated using these cells ^36^. Following infection, A549-hACE2 cells had significant attenuation of the YF mutant as compared to control (**Fig. 3F**). While no detectable virus was observed at 24 hours in either sample, at both 48 and 72 hours, the WT virus far exceeded the mutant in terms of viral load. These results confirmed attenuation of the YF mutant and argue that SG responses drive differences in viral load following infection.

### SARS-CoV-2 YF mutant attenuates replication *in vivo*

We next evaluated whether loss of FXR1 binding also attenuates viral replication *in vivo* using the golden Syrian hamster model for infection (**Fig. 4A**)^37–39^. Briefly, three-to-four-week old male hamsters were intranasally challenged with 10^5^ FFU of either SARS-CoV-2 WT or YF (**Fig. 4A**) and monitored for weight loss and disease over 7 days. At 2-, 4-, and 7-days post-infection (dpi), animals were nasal washed and euthanized for tissue collection to determine viral load changes. Similar to our previous work^30^, YF infected hamsters exhibited similar weight loss and disease to the WT cohort (**Fig. 4B**). However, viral load in the upper and lower respiratory tract revealed significant differences. On days 2 and 4 dpi, viral titers from nasal washes were lower in YF infected animals compared to WT (**Fig. 4C-D**). On day 4 pi, YF infected hamsters displayed significantly lower titers compared to WT infected group (**Fig. 4C-D**). No virus was detectable at 7 dpi in both specimens. These results reflect our *in vitro* findings and show that disrupting FXR1 binding with two residues in the NSP3-HVR are critical for attenuating replication.

**Figure 4:**
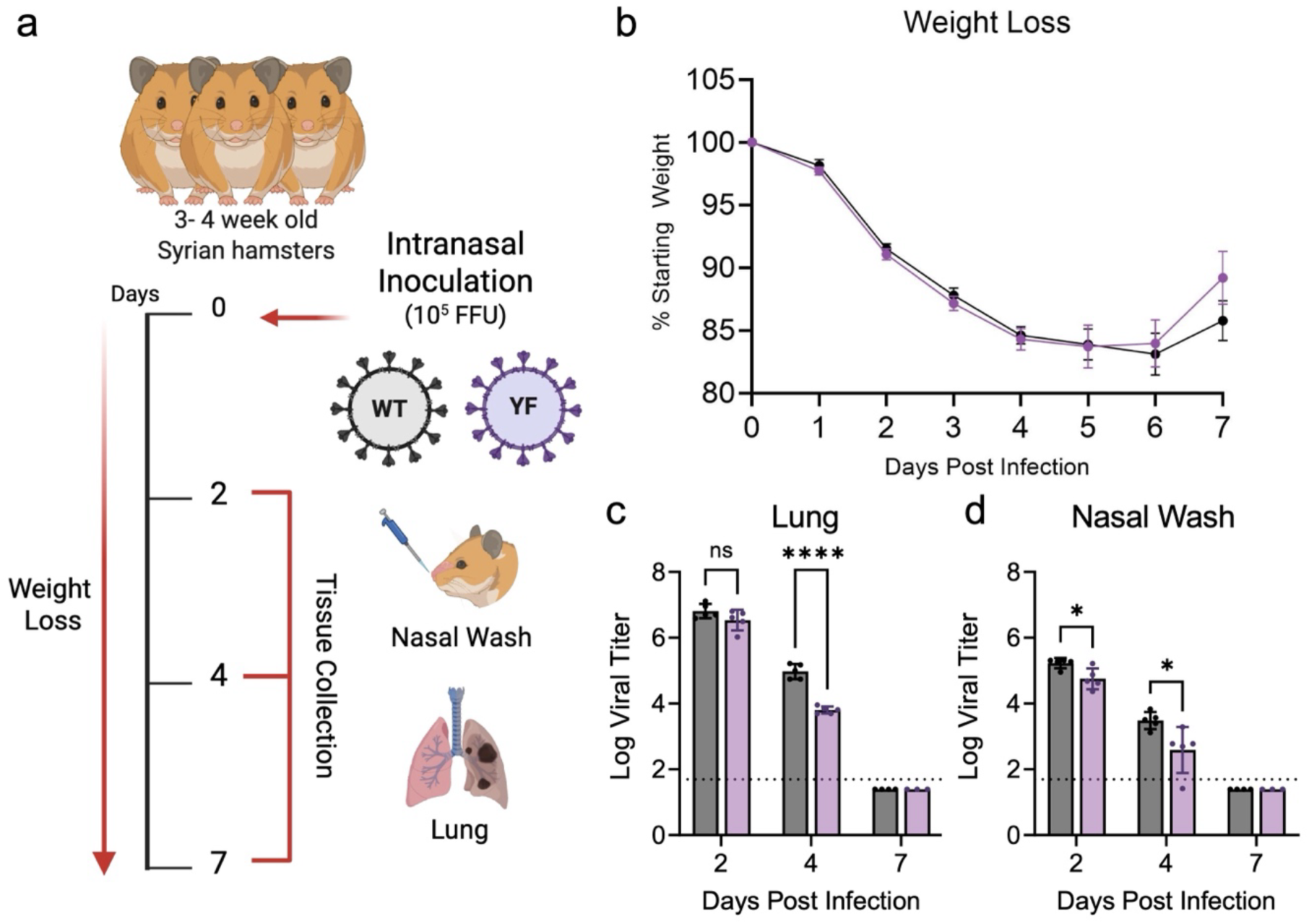
YF mutant has modest attenuation *in vivo*. (A) Schematic of golden Syrian hamster infection with WA-1 (WT) or YF mutant (purple) made using Biorender. Three-to four-week-old male hamsters were intranasally inoculated with 10^5^ FFU of WT or YF and monitored daily for signs of disease and weight loss (B) over 7 days. Infectious titers were measured in the lungs (C) and nasal wash (D) on 2-,4-, and 7-DPI. Each dot represents an infected animal. Statistical significance was determined by two-tailed Student’s t test. The dotted lines represent the assay limit of detection. Statistical significance was determined by two-tailed Student’s t test. *P ≤ 0.05; **P ≤ 0.01; ***P ≤ 0.001, ****p<0.0001. NS signifies not significant.

### Attenuation of YF mutant is not due to augmented interferon responses

Stress granule induction and type I interferon responses are both antiviral responses induced by the cell ^40–42^. While NSP3 binding to FXR1 links to stress granule formation, its implication for type I interferon and related responses are less clear. Here, we evaluated if the YF mutation sensitizes SARS-CoV-2 to type I interferons (IFN) responses. A key difference amongst the cell lines used are that Calu3 and A549 cells are type I IFN competent.^33^ Vero E6 cells cannot produce type I IFN but can respond to exogenous treatment^43^. Therefore, we investigated the effect of type I IFN on YF mutant replication. We pretreated Vero E6 cells with 100 U of recombinant of type I IFN (IFN-α) 16 hours prior to infection and compared samples against mock (PBS) pretreated controls. Similar to previous work, WT virus had only modest sensitivity to type I IFN pretreatment and resulted in a modest ∼ log reduction in viral titer at 24 hours (**Fig. 5A**). Similarly, the YF mutant also had modest sensitivity to type I IFN pretreatment; viral titer reduced ∼ 1-log at 24 HPI. In contrast to type I IFN sensitive SARS-CoV-2 mutants that show >3 log attenuation following IFN pretreatment ^44^, the similarity in reduction between the WT and YF mutant indicates that type I IFN responses are not the main driver of attenuation.

**Figure 5:**
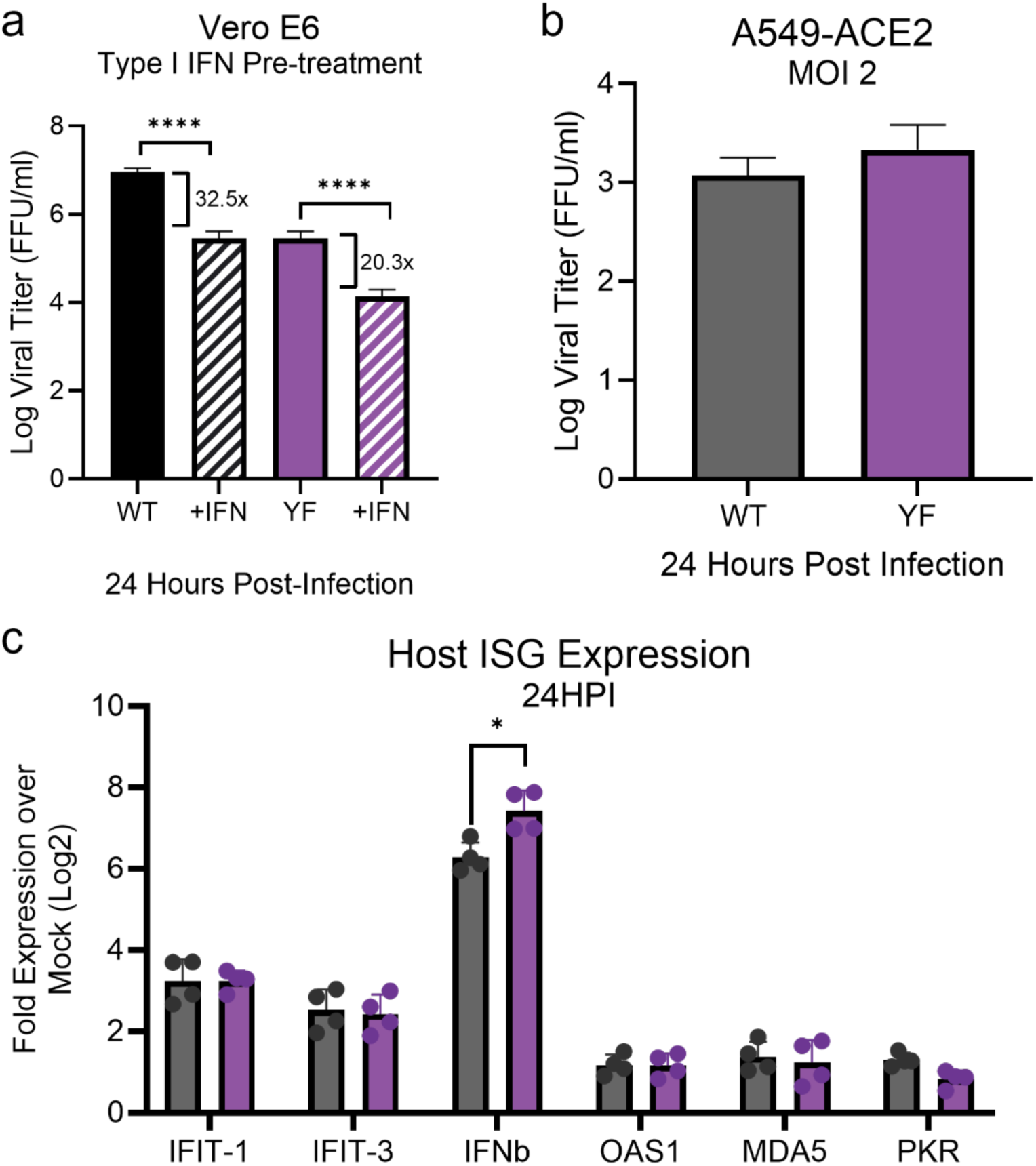
Decreased YF mutant replication is not driven by innate immune responses. (A) Vero E6 cells were pretreated with 100 U of recombinant type 1 IFN (hashed bar) or mock (solid bar) for 16 hours prior to infection. Cells were subsequently infected with either SARS-CoV-2 WT (black) or YF mutant (purple) at an MOI of 0.01. Viral titers were measured at 24 HPI. The fold change relative to mock control is shown. (B) A549-ACE2 cells were mock, WT, or YF infected at an MOI 1 for 24 hours. Viral titers at 24 HPI are shown. At 24 HPI, total RNA was collected and host mRNA expression was quantified by RT-qPCR. C_t_ values were normalized to human GAPDH and expressed as fold change (log_2_) of expression of indicated genes. Each dot represents a technical replicate, with the means for each target displayed, ± SD (error bars). Statistical analysis measured by two-tailed Student’s t test. *P ≤ 0.05; **P ≤ 0.01; ***P ≤ 0.001, ****p<0.0001.

To further examine if the attenuation of the YF mutant is driven by host innate immune pathways, we investigated the induction of type I and interferon stimulated genes. The largest differences between WT and YF mutant in viral titer were observed in A549-ACE2 cells which have intact SG and type I IFN responses (**Fig. 3F**)^45,46^. Here, we sought to equalize the viral load by infecting at a high multiplicity of infection (MOI 2) to ensure maximum infection and compare host expression differences between WT and mutant virus. Following infection, we found that viral replication was equivalent between WT and YF mutant 24HPI (**Fig. 5B**). We subsequently examined expression of innate immune response genes including type I IFN (*IFNB1*) and several IFN stimulated genes (*IFIT1, MDA5, OAS1b, PKR, IFIT3,)* (**Fig. 5C)**. While we note a modest increase in *IFNB1* expression in the mutant, the other ISGs are largely similar suggesting differential innate immune responses are not responsible for attenuation of the YF mutant.

### Stress granule formation is observed in YF mutant infected cell lines early during infection

Our prior studies demonstrated that SARS-CoV-2 NSP3 disrupts incorporation of FXR1 into SGs by competing with UBAP2L in a virus-free protein expression system ^30^. However, it is unclear if SG induction occurs in the context of infection with the SARS-CoV-2 YF mutant. Complicating this analysis, the N protein of SARS-CoV-2 has been shown by our group to be a potent SG inhibitor, though we predict its effect occurs primarily at late time points ^20^. Therefore, to evaluate SG dynamics following WT and YF mutant infection, we infected A549-ACE2 at 4, 6, and 10 HPI at an MOI of 1 and analyzed SGs by immunofluorescence. Visualizing infected cells, we stained for SARS-CoV-2 nucleocapsid (red), G3BP1 (green), and nuclei (Hoechst) and observed varying motifs at 6HPI (**Fig. 6A-C**). We first examined the percentage of cells that had the formation of G3BP1 foci and found no significant induction in any of the conditions at 4 HPI (**Fig. 6D**). However, by 6 hpi, we observed that ∼11% of cells in the YF infected group formed at least one SG as measured by G3BP1 foci (**Fig. 6C-D**). In contrast, <1% of cells were observed to have G3BP1 foci in WT and mock infected cells at 6HPI (**Fig. 6A-B & D**). Notably, we found low nucleocapsid staining in WT, but N appears to aggregate at the SG in the YF mutant as seen in the merge (**6C, white arrows**). These results correspond with SG levels peaking at 6HPI in YF infection, but by 10HPI, SG levels had reduced significantly to 2.7% in the YF mutant, although still elevated relative to 0.6% in WT. These results indicate that N protein has a potent impact on SGs at later times post infection.

**Figure 6:**
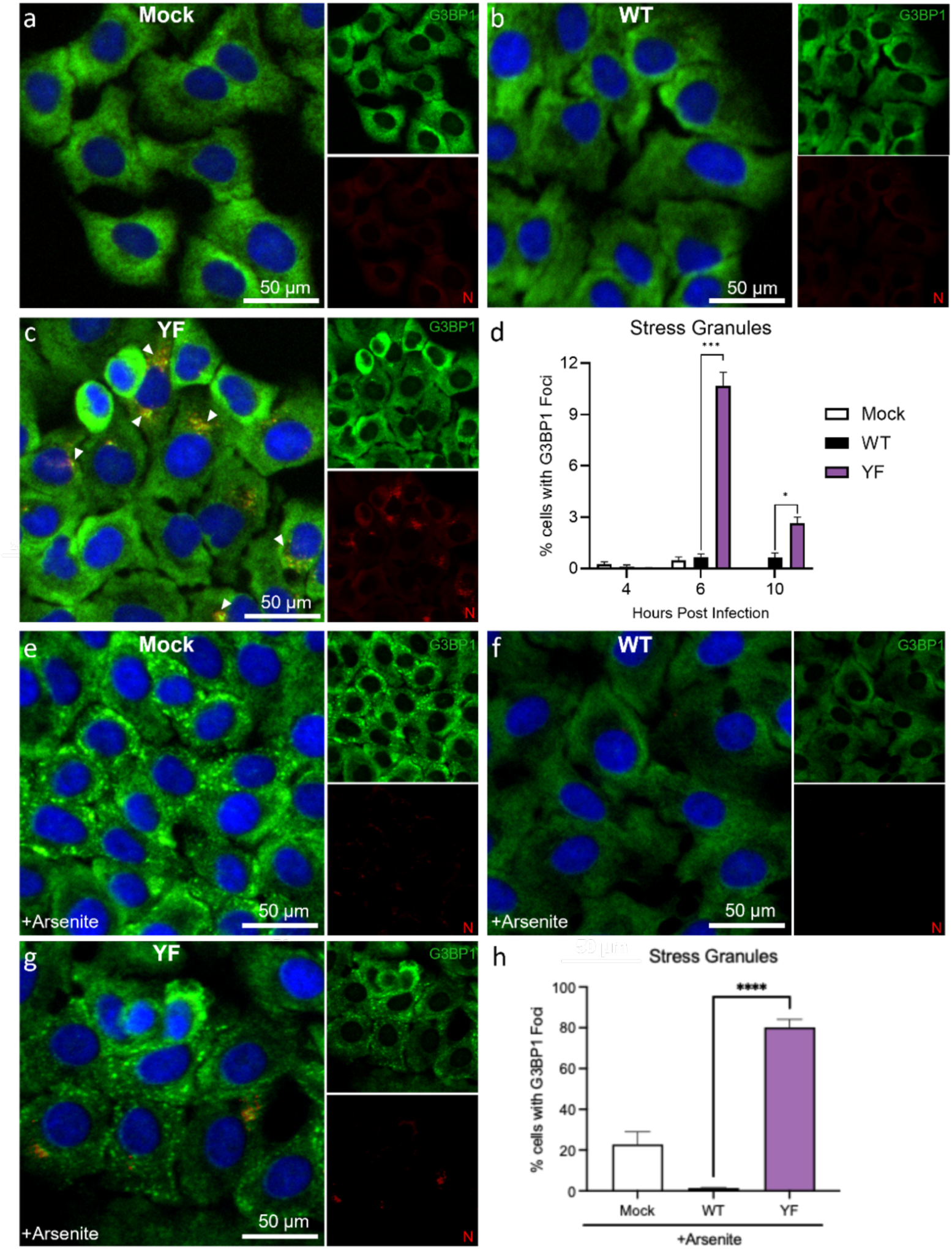
Stress granule formation is restored in YF mutant infected A549-ACE2 cells during early infection. A549-ACE2 cells were infected with WT (B) or YF mutant (C) at an MOI of 1 for 4, 6, and 10 HPI. Cells treated with PBS (A)were used as a control. G3BP1 (green), Nucleocapsid (red), and nuclei (blue) were labeled to visualize by immunofluorescence imaging. (A-C) Images at 6HPI are represented. (D) The total percent of cells with G3BP1+ foci (indicates SGs) were calculated using Fiji (ImageJ) software. (E-H) A549-ACE2 cells were infected WT (F) or YF mutant (G) at an MOI of 1 for 6 HPI. Cells treated with PBS (E) were used as a control. Thirty minutes prior to 6 hr timepoint, cells were treated with 1mM sodium arsenite to induce SG formation. (H) The total percent of cells with G3BP1+ foci at 6 HPI were calculated All quantitative data are shown as mean ± SD. Statistical analysis measured by two-tailed Student’s t test. *P ≤ 0.05; **P ≤ 0.01; ***P ≤ 0.001, ****p<0.0001.

Given the low amount of SG formation in standard infection (∼10%), we next determined if the YF mutant could inhibit SG formation if actively induced. To test this possibility, we treated cells with sodium arsenite, a chemical inducer of oxidative stress and SG formation. Mock cells treated with sodium arsenite induced SGs as measured by punctate G3BP1 foci (**Fig. 6E & H**). In contrast, WT infected cells actively inhibited SG formation and showed no evidence of G3BP1 foci (**Fig. 6F & H**). Importantly, YF infected cells had roughly 80% of cell showing G3BP1 foci (**Fig. 6G & H**). These results demonstrate that the YF mutant not only fails to inhibit SG formation at the early stage of infection but also augments SG formation beyond arsenite treatment alone. Overall, our results indicate that the loss of NSP3/FXR1 binding in the YF mutant induces a significant increase in SG positive cells by 6 HPI. While these dissipate at late times, likely due to N activity, it highlights the role of NSP3 and binding to FXR1 in disrupting SG formations early following infection.

## Discussion

Our previous studies revealed a functional role for the SARS-CoV-2 HVR in NSP3 for binding FXR1 and disrupting SG processes. Here, we extended this work by identifying residues Y138 and F145 in the NSP3 HVR as required for binding with FXR1. Subsequently mutating these residues (Y138A/F145A), we demonstrated that the SARS-CoV-2 YF mutant had attenuated replication *in vitro* relative to WT. While the YF mutant had similar disease to WT SARS-CoV-2 *in vivo*, the viral loads in the upper and lower respiratory tract were significantly decreased. Importantly, we showed that attenuation of the YF mutant is not due to changes in type I IFN sensitivity. Rather, reduced viral replication corresponded to the loss of SG control at early time points post infection. Overall, our study identifies Y138 and F145 within the NSP3-HVR as critical residues necessary for SARS-CoV-2 inhibition of stress granule formation at the early times post infection.

Stress granule formation is a highly dynamic process and has important implications for host cell functions. In the context of antiviral defense, the contribution of SGs are complex as both translation arrest and induction of innate immune signaling critically impact viral infection. Yet, the strongest evidence for the antiviral role of SGs is the genetic capital viruses use to limit its activation. In the case of SARS-CoV-2, the virus has two distinct viral proteins that contribute to control of SGs: nucleocapsid and NSP3 (**Fig. 7**). Our group and others have shown that nucleocapsid of SARS-CoV-2 interacts with G3BP1 to impair SG formation and inhibit innate immune signaling^20,47^. Similarly, our group has previously shown that the HVR of NSP3 binds to FXR1 and disrupts SG formation during the early phase of infection by competing with UBAP2L ^30^. Here, we define residues Y138 and F145, large aromatic amino acids in the HVR, are key to FXR1 binding and disrupting interactions with UBAP2L (**Fig. 1 & 2**). Importantly, these residues are completely conserved in the NSP3 HVR of the surveyed Sarbecoviruses (**Fig. 3B**) and likely indicate that stress granule antagonism is conserved across the family. Notably, while other CoV families do not have a similar motif in their NSP3 HVR, different CoV proteins have been shown to also antagonize SG function including MERS-CoV ORF4a, HCoV-OC43 NSP1, SARS-CoV-2 NSP5, and IBV NSP15 ^48–51^. Together, our results highlight the importance of controlling SG responses during CoV infection. In addition, it indicates that CoVs employ multiple mechanisms to limit stress granule activity.

**Figure 7:**
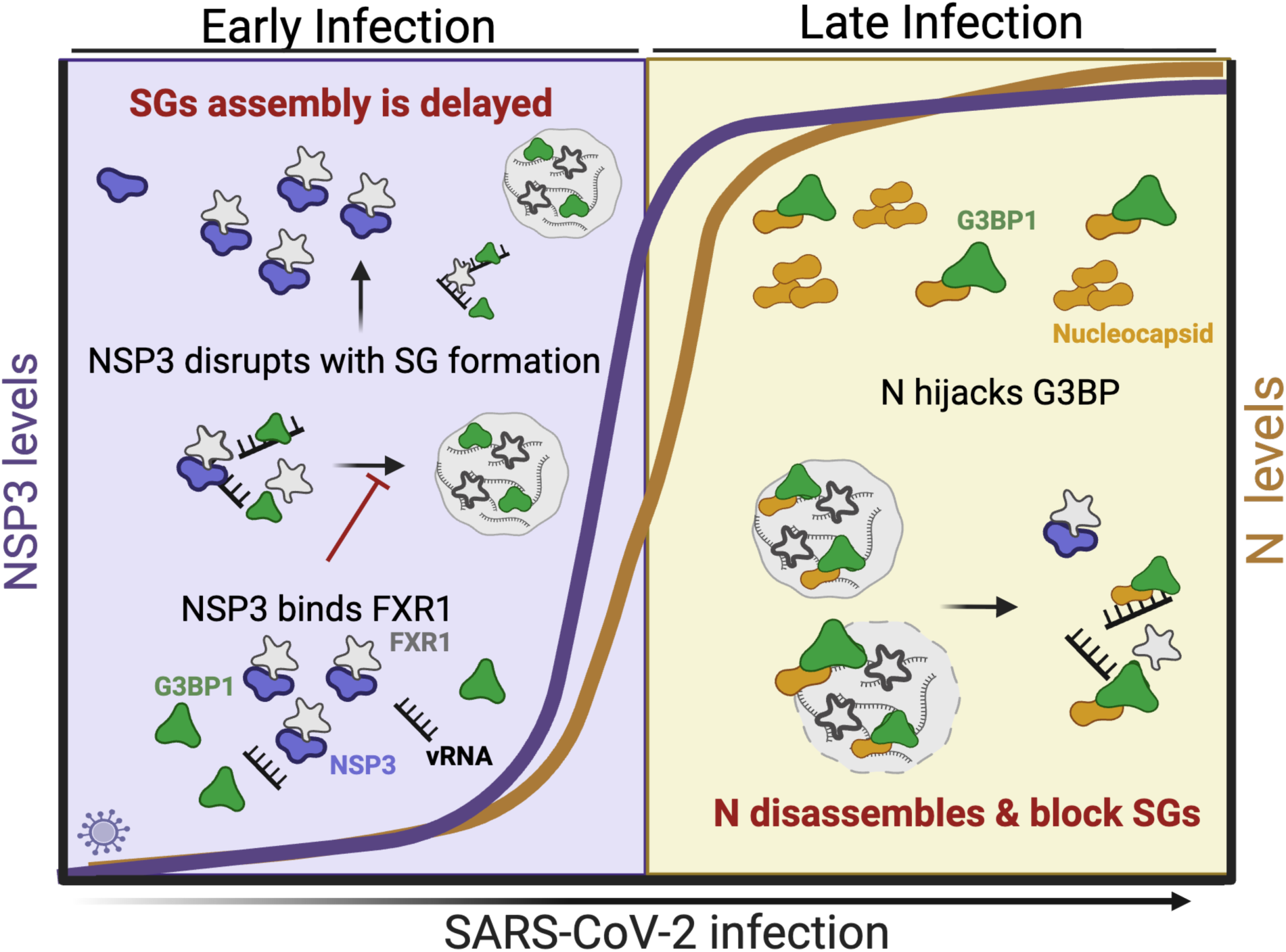
Early vs late SG regulation. Proposed model of SARS-CoV-2 NSP3 role in SG regulation at early stages of infection. (A) During early phase of infection, NSP3 HVR binds to FXR1 to disrupt and delay SG formation. At the late stage of infection, SG formation is inhibited by N protein when expression levels are highest. Overall, NSP3 and N protein work in cooperation to kinetically regulate SG formation during the course of infection.

Stress granule formation is an evolutionary conserved process found in both mammals and plants to combat stress conditions ^9,52^. As such, viruses have also evolved dedicated processes to prevent SG formation. Here, we have shown that two FXR1 binding residues in NSP3 are conserved in SARS-CoV and bat related Sarbecoviruses. Other work has shown that Y138 is not conserved in MERS, hCoV-229E, or hCoV-OC43^21^; however, F145 is conserved in 229E and OC43 suggesting potential conservation of theses amino acids. Specifically, to act as a macromolecular hub for protein assembly needed for efficient viral replication^21^. Aromatic amino acids such as phenylalanine and tyrosine have been shown to be crucial in mediating LLPS by driving multivalent interactions.^53,54^ Therefore, we cannot exclude that these same HVR residues could have an impact on SG formation by other CoVs outside of SARS-CoV-2. Notably, antagonism of SGs by FXR1 is a unique finding. FXRs have been demonstrated to play pivotal roles in viral replication in several RNA viruses and especially defined during alphavirus infection. In the case of new world alphaviruses, VEEV and EEEV have adopted strategies to use FXRs to drive the assembly of viral replication complexes (vRCs) through interaction with their NSP3 hypervariable domain (HVD).^23,28,55^ Conversely, FXR proteins were recently demonstrated to drive double membrane vesicle (DMV) clustering through the same region of NSP3-HVR investigated here. ^21,56^ However, sequence alignment of old and new world alphavirus HVDs revealed there is no presence of conserved tyrosine of phenylalanine throughout the domain. Thus, targeting FXR1 to inhibit SGs by key residues may be a unique feature adopted by CoVs.

In recent years, the link between SG responses and host antiviral defenses have complicated by conflicting studies. Studies have argued that SGs amplify antiviral activity by serving as signaling platforms to promote the induction of IFN responses ^17,57^. Alternatively, other groups have described a role for SGs as “shock absorbers” to regulate over activation of innate immunity^58^. Others have indicated loss of SG function had no impact on innate immune responses^19^. In this study, we find that loss of early SG control by the YF mutant does not augment sensitivity to type I IFN (**Fig. 5A**). Similarly, minimal changes in type I IFN or ISGs were observed in the YF mutant as compared to control, suggesting SG induction has limited impact on innate immune signaling (**Fig. 5C**). However, accumulation of the N protein and its antagonism of SGs at later times complicates these findings. Importantly, the YF mutant attenuation is not driven by type I IFN responses but rather corresponds to loss of SG control at early times compared to WT (**Fig. 6**). Overall, the findings indicate activating SG responses may offer a novel route to limit viral infection and pathogenesis.

In addition to defining the key residues in NSP3 driving FXR1 binding, our study also establishes a kinetic profile for SG antagonism by SARS-CoV-2. The N protein has been shown to be a powerful antagonist of SGs by disrupting G3BP1 activity even after SGs have formed^47,59–62^. However, N has several roles during viral infection including genome packaging, viral assembly, regulation of viral transcription, and modulation of host factors ^63–66^. These roles are likely controlled by the relative abundance of the N protein and post-translational modifications like phosphorylation ^65,67^. Here, we predict that NSP3 and N act at different kinetic times to antagonize SG activity (**Fig. 7)**. We expect N activity against SG occurs later during infection corresponding to its peak production approximately at 10 HPI^68^. In this scenario, SARS-CoV-2 NSP3 functions to disrupt SG at earlier points during infection. At 4 HPI, we detect no difference in SG accumulation between our mutant and WT (**Fig. 6D**); however, by 6 HPI, we detected a higher fraction of SG positive cells in our YF infected group compared to control. This dynamic proposes a kinetic framework by which SARS-CoV-2 NSP3 and N work in cooperation to regulate the SG responses.

Overall, our manuscript identifies aromatic residues Y138 and F145 as critical to the binding of FXR1 and disruption of SG formation during the early parts of SARS-CoV-2 infection. The YF mutant that ablates NSP3 interactions with FXR1 has attenuated growth *in vitro* and *in vivo* that is not governed by augmented type I interferon responses. Instead, we find that the loss of the Y138/F145 results in augmented SG formation at early times and corresponds to attenuated viral replication. Importantly, in combination with the N proteins, our data highlights the genetic capital SARS-CoV-2 invests in controlling SG responses during the course of infection.

## Materials & Methods

### Cell Lines

African green monkey kidney Vero E6 cells were cultured in high glucose Dulbecco’s modified Eagle medium (DMEM)(Gibco #11965-092) supplemented with 10% Fetal Clone II (FBS)(Hyclone; #SH30071.03) and 1% antibiotic/antimycotic (anti-anti) (Gibco #5240062). Vero E6 expressing human TMPRSS2 were grown in high glucose DMEM supplemented with 10% FBS, 1% anti-anti, and 1 mg/mL Geneticin (G418) (Gibco; 10131035). Calu-3-2b4 (Calu-3) cells were grown in high glucose DMEM with 10% defined FBS, 1% anti-anti, and 1 mg/mL sodium pyruvate (1mM). A549 cells expressing human ACE2 were grown in 10% FBS, 1% anti-anti, and 10 µg/mL Blasticidin S HCL (Gibco; # A1113902).

Vero E6 and Calu-3 cells were provided by Dr. Ralph Barric. A549-ACE2 cells were provided by Dr. Pei Yong Shi. All cell lines grow at 37°C with 5% CO_2_ and are mycoplasma tested periodically. (Last negative test in 2024). Single cell FACS sorting (BD Aria Fusion) was applied on Vero-TMPRSS2 and A549-ACE2 cells to generate TMPRSS2 or ACE2 positive cell populations.

### Viruses

#### Reverse Genetic Clones

All recombinant viruses were generated using reverse genetics as previously described ^31,32^. WT and mutant SARS-CoV-2 sequences are used from the USA-WA1/2020 isolate sequence provided by the Word Reference Center for Emerging Viruses and Arboviruses (WRCEVA), originally obtained from the US Centers for Disease Control and Prevention (CDC) as previously described ^69,70^. The NSP3 YF mutant was constructed using standard cloning techniques as previously outline for our reverse genetics system. ^31,32^ with virus stocks being amplified on Vero E6-TMPRSS2 to prevent mutations from occurring in furin cleavage site (FCS). ^71^ All viruses were expanded once (P1) for use in studies and tittered by Focus-forming assay (FFA) and plaque assay (for plaque morphology). All infections and manipulations were performed in a biosafety level 3 (BSL-3) laboratory in accordance with approved protocols and use of appropriate personal protective equipment. P1 viral stocks were verified through Sanger sequencing of cDNA for mutations in spike furin cleavage site and the NSP3-HVR for the introduced mutations.

### Biosafety

The synthetic construction of SARS-CoV-2 NSP3 mutants was reviewed for DURC/P3CO policies and approved by the University of Texas Medical Branch Institutional Biosafety Committee. All studies in animals were conducted under a protocol approved by the UTMB Institutional Animal Care and Use Committee and complied with USDA guidelines in a laboratory accredited by the Association for Assessment and Accreditation of Laboratory Animal Care. UTMB is a registered Research Facility under the Animal Welfare Act. It has a current assurance (A3314-01) with the Office of Laboratory Animal Welfare (OLAW), in compliance with NIH Policy. Procedures involving infectious SARS-CoV-2 were performed in the Galveston National Laboratory ABSL3 facility.

### *In vitro* infection

#### Viral replication kinetics

Viral replication studies were performed as previously described.^30^ Briefly, cells (Vero E6, A549-ACE2) were seeded in seeded in six well plates the day before infection. Calu-3 cells were seeded at high density and allowed to growth to confluency in six wells plates before conducting growth curve. Cells were infected at indicated MOI for 45 minutes at 37°C with 5% CO_2_ and every 15 minutes. After absorption, plates were washed three times with PBS, and fresh media was replaced in wells, indicating time zero. Wells were sampled at specified time points. Three technical replicates were collected for each timepoint, and each experiment was performed twice. Samples were titrated with focus-forming assay.

For high MOI infections, A549-ACE2 cells were plated in 12-well plates at 5×10^5^ cells per well. Viruses were diluted in PBS and added to cells and incubated for 45 minutes at 37°C with 5% CO_2._ and rocked every 15 minutes. Cells were washed three times and replaced with complete DMEM (10% FBS with 1x Anti-Anti). For each time point, 200 ul of supernatant was collected for viral titers and cells were lysed for qPCR gene expression analysis with Trizol ™.

#### Type I IFN Assay

For IFN pretreatment, 100 Units (U)/mL of recombinant IFN-β (PBL Laboratories) was added to Vero E6 cells 16-18 hours prior to infection. Growth curves were conducted as mentioned above.

### *In vivo* infection

#### Animal studies

Three-to four-week male Syrian golden hamsters (HsdHAN:AURA strain) were purchased from Envigo, Indianapolis, IN. For infection, hamsters were intranasally inoculated with 1×10^5^ FFU of SARS-CoV-2 WT or YF mutant in a 100 µL volume. Infected animals were weighed and monitored daily for illness over 7 days. Cohorts of 5 animals per group (including mock infected animals) were anesthetized with isoflurane and nasal washes were collected in 400 µl PBS on endpoint days (2, 4, and 7 DPI) for titers. Animals were then humanely euthanized by CO_2_ immediately following nasal washes for lung specimen collection. Lobes were stored in RNAlater solution (Invitrogen) for qPCR analysis and in PBS for titer analysis.

### Focus-forming assay

Focus-forming assays (FFAs) were performed as previously described with adaptations. ^72^ Briefly, Vero E6 cells were seeded in 96-well black walled plates to be 100% confluent the next day. Cell culture supernatants, nasal washes, or homogenized tissues containing SARS-CoV-2 underwent 10-fold serial dilutions in serum-free media. Twenty µl of diluted and undilute sample were added to wells and incubated for 45 minutes with rocking every 15 minutes. After absorption, methylcellulose overlay (0.85%) was then added to each well and cells were incubated for 24 hours. Next day, the methylcellulose was removed, and cells were washed three times with PBS and fixed in 10% buffered formalin for 30 minutes at room temperature (rt) to inactivate virus. For staining, cells were permeabilized and blocked with PBS solution containing 0.1% saponin and 0.1% BSA (Perm. Buffer) for 30 minutes. Following incubation, α-SARS-CoV-2 Nucleocapsid primary antibody (Cell Signaling, 68344) at 1:3,000 in Perm. buffer overnight at 4◦C. Cells were washed 3 times with 1X DPBS before incubating with Alexa Fluor 555-conjugated α-mouse secondary antibody (Invitrogen, A28180) at 1:2,000 in Perm. Buffer for 1 hour at rt wrapped in foil. Cells were washed 3 times with 1X DPBS and fluorescent foci images were captured using a Cytation 7 cell imaging multi-mode reader (Biotek). Foci were counted using ImageJ free software (NIH).

#### Homogenization

Hamster lungs were infected as described in ‘Animal Studies’ and the right inferior lobe were stored in sterile PBS, zirconia beads, and kept at −80◦C. Tissues were homogenized using a MagNALyser (Roche Life Science) at 6,000 rpm for 1 minute total.

### Real-Time quantitative PCR (qPCR)

#### RNA Extraction & qPCR

Total RNA from cells were isolated with Direct-zol RNA Miniprep Plus kit (Zymo, #R2072) according to manufacturer’s protocol. RNA was reverse transcribed to complimentary DNA (cDNA) using the iScript cDNA synthesis kit according to manufacturer’s instructions (Bio-Rad, 1708891). RT-qPCR was performed on cDNA using the Luna Universal qPCR Master Mix (NEB #M3003) per manufacturer’s instructions. Fluorescent readings were measured on a Bio-Rad CFX Connect instrument using Bio-Rad CFX Maestro 1.1 (version 4.1.2433.1219). The primers used to amplify host targets were human GAPDH (forward:5’-TCAAGATCATCAGCAATGCC; reverse: 5’- AAGTTGTCATGGATGACCTTGG), IFIT1 (forward:5’-CCAAGGAGACCCCAGAAACC; reverse: 5’-CGCTACGTGGAGTGAGCTAG), IFIT3 (forward:5’-AAGAACAAATCAGCCTGGTCAC; reverse: 5’-TCCCTTGAGACACTGTCTTCC), IFNβ (forward:5’-AGTAGGCGACACTGTTCGTG; reverse: 5’-AGCCTCCCATTCAATTGCCA), OAS1 (forward:5’-GAGCTCCTGACGGTCTATGC; reverse: 5’-TCATCGTCTGCACTGTTGCT), MDA5 (forward:5’-AAGCCCACCATCTGATTGGAG; reverse: 5’-CCACTGTGGTAGCGATAAGCAG),PKR (forward:5’- GAAGTGGACCTCTACGCTTTGG; reverse: 5’- GATGATGCCATCCCGTAGGTC). (Average Ct values of technical replicates were normalized using the ΔΔCt method to the house keeping gene (human GAPDH) and expressed as Log_2_ relative fold change.

### Immunofluorescence and microscopy

A549-ACE2 cells were seeded on 96-well black sided plates one day prior to infection. Infections were conducted at indicated MOI in an inoculum volume of 50 µL volume per well. Incubation was conducted as mentioned in ‘viral replication kinetics. Wells were replaced with fresh cell culture media containing no selective antibiotics. To induce SG formation, wells were replaced with cell culture media containing 1mM sodium arsenite 30 minutes prior to inactivation timepoint. Plates were inactivated in 10% buffered formalin at indicated HPI. Antibody staining was adapted from Lamichhane et. Al, 2024. Briefly, fixed cells were permeabilized with 0.2% Triton X-100 in PBS for 10 minutes at rt and blocked with 2% BSA for 1 hour. Samples were then incubated with primary antibodies; Rabbit anti-G3BP1 (Cell signaling #61559) and mouse anti-Nucleocapsid (Cell Signaling, 68344BC) overnight at 4◦C. The next day, cells were washed 3 times with PBS and incubated with host-specific Alexa Fluro 488 (Invitrogen; A32731) and Alexa Fluro 555 (Invitrogen; A28180) secondary antibodies for 1 hour at rt, away from light.^73^ After secondary incubation, plates were washed 3 times, and nuclei were stained with 1 ug/ml Hoecsht 3342 (Thermo Fisher, H1399) according to manufacturer instructions for 15 minutes at rt. Images were captured using a Cytation 7 instrument at 20x magnification.

### Stress granule quantification

Quantification of SGs was adapted from Dolliver et. Al., 2022. Briefly, the total percent of G3BP1 positive cells were performed by counting the total number of cells based on Hoechst staining. First, the total number of nuclei were counted using a self-developed Image J macro, to automatically threshold and count total cells in an area. At least four areas were imaged in one well (repeated in triplicate) with consistent exposure setting for all images. Cells containing at least 1 granule we counted as SG+ using the multi-point tool in ImageJ.^48^ A minimum of 50 cells per area were counted for each condition in each experiment. Images were analyzed using ImageJ 1.54p Fiji. For robust image analysis, standardized image acquisition practices were followed. Low to moderate cell densities were imaged to prevent negative impacts on cell counting and segmentation. Image acquisition was carried out using optimal constant exposure settings for each experiment. Background signal subtraction was performed using negative controls in ImageJ.”The QuickFigures plugin for ImageJ was used to prepare immunofluorescence panels. ^74^

### Structure modeling

Structural models were generated to visualize FXR1 and NSP3 residue interactions. using AlphaFold as previously described. ^30^ Models were generated using PyMOL (version 3.10.) for WA-1 NSP3 and human FXR1. Residues Y138 and F145 are highlighted and mutated to visualize alanine mutations in YF residues of NSP3.

### Immunoprecipitations

HeLa cells were transfected with with 2 μg DNA and 2μL Jet Optimus reagent overnight. Cell pellets were collected after 24 hours and lysed in 400 μL lysis buffer (100mM NaCl, 50mM Tris pH 7,4, 0,1% NP40, 0,2% Triton-100, 1mM DTT) supplemented with protease (complete mini EDTA free, Roche) and phosphatase inhibitor tablets (Roche) for 45 minutes on ice. Lysates were cleared at 20000g and 4°C for 1h. Cleared lysates were incubated with 10 μL pre-equilibrated GFP-trap beads for 1h at 4°C on a rotor-wheel. Following 3 washes with 1 mL wash buffer (150mM NaCl, 50mM Tris pH 7,4, 0,05% NP40, 5% Glycerol, 1mM DTT), the samples were eluted with 25 uL 2x LSB (Thermo) and processed for western blot using the following antibodies: FXR1 mouse (Santa Cruz 374148) and GFP rabbit (made in house against GFP).

### Isothermal calorimetry

ITC experiments were set up as in^30^. The following peptides were purchased from Peptide 2.0 Inc (Chantilly. VA, USA) were used:

NSP3 WT: QYEYGTEDDYQGKPLEFGATSW
NSP3 YF/AA: QYEYGTEDD**<u>A</u>**QGKPLE**<u>A</u>**GATSW

## Acknowledgements

Research was supported by grants from NIAID of the United States NIH (R01-AI153602, R21-AI145400, U19-AI171413) to VDM. The research was also supported by STARs Award provided by the University of Texas System and Burroughs Welcome Fund Investigators in Pathogenesis grant to VDM. Trainee funding provided by NIAID of the NIH to REA (T32AI007526-24). Work at the Novo Nordisk Foundation Center for Protein Research is supported by grant NNF14CC0001 and JN is supported by a grant from Sygeforsikring Danmark and JN by a grant from Independent Research Fund Denmark (3101-00358B). We thank Parimal, Megan, and Adam Hage for technical assistance on the manuscript.

## Competing Interests

VDM has filed a patent on the reverse genetic system for SARS-CoV-2. All other authors declare no conflicts of interest.

## Author Contributions

Conceptualization: REA, BAJ, JN, VDM

Formal analysis: REA, KGL, DG, BAJ, JN, VDM

Funding acquisition: REA, JN, VDM

Investigation: REA, KGL, DG, LKE, YA, AMM, AM, JC, BM, JAP, KSP

Methodology: REA, KGL, BAJ, VDM

Project Administration: BAJ, JN, VDM

Supervision: BAJ, JN, VDM

Visualization: REA, VDM

Writing – original draft: REA, VDM

Writing – review and editing: REA, BAJ, VDM

## REFERENCES

1. Msemburi W, Karlinsky A, Knutson V, Aleshin-Guendel S, Chatterji S, Wakefield J. The WHO estimates of excess mortality associated with the COVID-19 pandemic. Nature. 2023;613(7942):130–7.

2. Fehr AR, Perlman S. Coronaviruses, Methods and Protocols. Coronaviruses. 2015;1282:1–23.

3. Perlman S, Netland J. Coronaviruses post-SARS: update on replication and pathogenesis. Nat Rev Microbiol. 2009;7(6):439–50.

4. Báez-Santos YM, John SESt, Mesecar AD. The SARS-coronavirus papain-like protease: Structure, function and inhibition by designed antiviral compounds. Antivir Res. 2015;115:21–38.

5. Wang Y, Grunewald M, Perlman S. Coronaviruses: An Updated Overview of Their Replication and Pathogenesis. Methods Mol Biol. 2020;2203:1–29.

6. Lei J, Kusov Y, Hilgenfeld R. Nsp3 of coronaviruses: Structures and functions of a large multi-domain protein. Antivir Res. 2018;149:58–74.

7. Shi M, Wu Z, Zhang Y, Li T. Decoding intrinsically disordered regions in biomolecular condensates. Fundam Res. 2025;

8. Guan Y, Wang Y, Fu X, Bai G, Li X, Mao J, et al. Multiple functions of stress granules in viral infection at a glance. Front Microbiol. 2023;14:1138864.

9. Protter DSW, Parker R. Principles and Properties of Stress Granules. Trends Cell Biol. 2016;26(9):668–79.

10. Tourrière H, Chebli K, Zekri L, Courselaud B, Blanchard JM, Bertrand E, et al. The RasGAP-associated endoribonuclease G3BP mediates stress granule assembly. J Cell Biol. 2023;222(11):e200212128072023new.

11. Hofmann S, Kedersha N, Anderson P, Ivanov P. Molecular mechanisms of stress granule assembly and disassembly. Biochim Biophys Acta (BBA) - Mol Cell Res. 2021;1868(1):118876.

12. Siomi MC, Siomi H, Sauer WH, Srinivasan S, Nussbaum RL, Dreyfuss G. FXR1, an autosomal homolog of the fragile X mental retardation gene. EMBO J. 1995;14(11):2401–8.

13. Kirkpatrick LL, McIlwain KA, Nelson DL. Comparative Genomic Sequence Analysis of the FXR Gene Family: FMR1, FXR1, and FXR2. Genomics. 2001;78(3):169–77.

14. Walsh D, Mohr I. Viral subversion of the host protein synthesis machinery. Nat Rev Microbiol. 2011;9(12):860–75.

15. Dhaka P, Singh A, Nehul S, Choudhary S, Panda PK, Sharma GK, et al. Disruption of Molecular Interactions between the G3BP1 Stress Granule Host Protein and the Nucleocapsid (NTD-N) Protein Impedes SARS-CoV-2 Virus Replication. Biochemistry. 2025;64(4):823–40.

16. Yang P, Mathieu C, Kolaitis RM, Zhang P, Messing J, Yurtsever U, et al. G3BP1 Is a Tunable Switch that Triggers Phase Separation to Assemble Stress Granules. Cell. 2020;181(2):325–345.e28.

17. Yang W, Ru Y, Ren J, Bai J, Wei J, Fu S, et al. G3BP1 inhibits RNA virus replication by positively regulating RIG-I-mediated cellular antiviral response. Cell Death Dis. 2019;10(12):946.

18. Hosmillo M, Lu J, McAllaster MR, Eaglesham JB, Wang X, Emmott E, et al. Noroviruses subvert the core stress granule component G3BP1 to promote viral VPg-dependent translation. eLife. 2019;8:e46681.

19. Burke JM, Ratnayake OC, Watkins JM, Perera R, Parker R. G3BP1-dependent condensation of translationally inactive viral RNAs antagonizes infection. Sci Adv. 2024;10(5):eadk8152.

20. Yang Z, Johnson BA, Meliopoulos VA, Ju X, Zhang P, Hughes MP, et al. Interaction between host G3BP and viral nucleocapsid protein regulates SARS-CoV-2 replication and pathogenicity. Cell Rep. 2024;43(3):113965.

21. Li M, Hou Y, Zhou Y, Yang Z, Zhao H, Jian T, et al. LLPS of FXR proteins drives replication organelle clustering for β-coronaviral proliferation. J Cell Biol. 2024;223(6):e202309140.

22. Hajikhezri Z, Kaira Y, Schubert E, Darweesh M, Svensson C, Akusjärvi G, et al. Fragile X-Related Protein FXR1 Controls Human Adenovirus Capsid mRNA Metabolism. J Virol. 2023;97(2):e01539–22.

23. Frolov I, Kim DY, Akhrymuk M, Mobley JA, Frolova EI. Hypervariable Domain of Eastern Equine Encephalitis Virus nsP3 Redundantly Utilizes Multiple Cellular Proteins for Replication Complex Assembly. J Virol. 2017;91(14).

24. Soto-Acosta R, Xie X, Shan C, Baker CK, Shi PY, Rossi SL, et al. Fragile X mental retardation protein is a Zika virus restriction factor that is antagonized by subgenomic flaviviral RNA. Elife. 2018;7:e39023.

25. Pereira B, Billaud M, Almeida R. RNA-Binding Proteins in Cancer: Old Players and New Actors. Trends Cancer. 2017;3(7):506–28.

26. Kang JY, Wen Z, Pan D, Zhang Y, Li Q, Zhong A, et al. LLPS of FXR1 drives spermiogenesis by activating translation of stored mRNAs. Science. 2022;377(6607):eabj6647.

27. Kim TH, Tsang B, Vernon RM, Sonenberg N, Kay LE, Forman-Kay JD. Phospho-dependent phase separation of FMRP and CAPRIN1 recapitulates regulation of translation and deadenylation. Science. 2019;365(6455):825–9.

28. Kim DY, Reynaud JM, Rasalouskaya A, Akhrymuk I, Mobley JA, Frolov I, et al. New World and Old World Alphaviruses Have Evolved to Exploit Different Components of Stress Granules, FXR and G3BP Proteins, for Assembly of Viral Replication Complexes. Plos Pathog. 2016;12(8):e1005810.

29. Foy NJ, Akhrymuk M, Akhrymuk I, Atasheva S, Bopda-Waffo A, Frolov I, et al. Hypervariable Domains of nsP3 Proteins of New World and Old World Alphaviruses Mediate Formation of Distinct, Virus-Specific Protein Complexes. J Virol. 2013;87(4):1997–2010.

30. Garvanska DH, Alvarado RE, Mundt FO, Lindqvist R, Duel JK, Coscia F, et al. The NSP3 protein of SARS-CoV-2 binds fragile X mental retardation proteins to disrupt UBAP2L interactions. EMBO Rep. 2024;1–25.

31. Xie X, Muruato A, Lokugamage KG, Narayanan K, Zhang X, Zou J, et al. An Infectious cDNA Clone of SARS-CoV-2. Cell Host Microbe. 2020;27(5):841–848.e3.

32. Xie X, Lokugamage KG, Zhang X, Vu MN, Muruato AE, Menachery VD, et al. Engineering SARS-CoV-2 using a reverse genetic system. Nat Protoc. 2021;16(3):1761–84.

33. Lokugamage KG, Hage A, Vries M de, Valero-Jimenez AM, Schindewolf C, Dittmann M, et al. Type I Interferon Susceptibility Distinguishes SARS-CoV-2 from SARS-CoV. J Virol. 2020;94(23):e01410–20.

34. Matsuyama S, Nao N, Shirato K, Kawase M, Saito S, Takayama I, et al. Enhanced isolation of SARS-CoV-2 by TMPRSS2-expressing cells. Proc Natl Acad Sci. 2020;117(13):7001–3.

35. Koch J, Uckeley ZM, Doldan P, Stanifer M, Boulant S, Lozach P. TMPRSS2 expression dictates the entry route used by SARS-CoV-2 to infect host cells. EMBO J. 2021;40(16):EMBJ2021107821.

36. Gottschald OR, Malec V, Krasteva G, Hasan D, Kamlah F, Herold S, et al. TIAR and TIA-1 mRNA-Binding Proteins Co-aggregate under Conditions of Rapid Oxygen Decline and Extreme Hypoxia and Suppress the HIF-1α Pathway. J Mol Cell Biol. 2010;2(6):345–56.

37. Chan JFW, Zhang AJ, Yuan S, Poon VKM, Chan CCS, Lee ACY, et al. Simulation of the clinical and pathological manifestations of Coronavirus Disease 2019 (COVID-19) in golden Syrian hamster model: implications for disease pathogenesis and transmissibility. Clin Infect Dis. 2020;71(9):ciaa325.

38. Bednash JS, Kagan VE, Englert JA, Farkas D, Tyurina YY, Tyurin VA, et al. Syrian hamsters as a model of lung injury with SARS-CoV-2 infection: Pathologic, physiologic, and detailed molecular profiling. Transl Res. 2022;240:1–16.

39. Imai M, Iwatsuki-Horimoto K, Hatta M, Loeber S, Halfmann PJ, Nakajima N, et al. Syrian hamsters as a small animal model for SARS-CoV-2 infection and countermeasure development. Proc National Acad Sci. 2020;117(28):16587–95.

40. Lei X, Dong X, Ma R, Wang W, Xiao X, Tian Z, et al. Activation and evasion of type I interferon responses by SARS-CoV-2. Nat Commun. 2020;11(1):3810.

41. Onomoto K, Yoneyama M, Fung G, Kato H, Fujita T. Antiviral innate immunity and stress granule responses. Trends Immunol. 2014;35(9):420–8.

42. Eiermann N, Haneke K, Sun Z, Stoecklin G, Ruggieri A. Dance with the Devil: Stress Granules and Signaling in Antiviral Responses. Viruses. 2020;12(9):984.

43. Diaz MO, Ziemin S, Beau MML, Pitha P, Smith SD, Chilcote RR, et al. Homozygous deletion of the alpha- and beta 1-interferon genes in human leukemia and derived cell lines. Proc Natl Acad Sci. 1988;85(14):5259–63.

44. Schindewolf C, Lokugamage K, Vu MN, Johnson BA, Scharton D, Plante JA, et al. SARS-CoV-2 Uses Nonstructural Protein 16 To Evade Restriction by IFIT1 and IFIT3. J Virol. 2023;97(2):e01532–22.

45. Murgolo N, Therien AG, Howell B, Klein D, Koeplinger K, Lieberman LA, et al. SARS-CoV-2 tropism, entry, replication, and propagation: Considerations for drug discovery and development. PLoS Pathog. 2021;17(2):e1009225.

46. Lujan H, Criscitiello MF, Hering AS, Sayes CM. Refining In Vitro Toxicity Models: Comparing Baseline Characteristics of Lung Cell Types. Toxicol Sci. 2019;168(2):302–14.

47. He S, Gou H, Zhou Y, Wu C, Ren X, Wu X, et al. The SARS-CoV-2 nucleocapsid protein suppresses innate immunity by remodeling stress granules to atypical foci. FASEB J. 2023;37(12):e23269.

48. Dolliver SM, Kleer M, Bui-Marinos MP, Ying S, Corcoran JA, Khaperskyy DA. Nsp1 proteins of human coronaviruses HCoV-OC43 and SARS-CoV2 inhibit stress granule formation. Plos Pathog. 2022;18(12):e1011041.

49. Gao B, Gong X, Fang S, Weng W, Wang H, Chu H, et al. Inhibition of anti-viral stress granule formation by coronavirus endoribonuclease nsp15 ensures efficient virus replication. Plos Pathog. 2021;17(2):e1008690.

50. Zheng Y, Deng J, Han L, Zhuang MW, Xu Y, Zhang J, et al. SARS-CoV-2 NSP5 and N protein counteract the RIG-I signaling pathway by suppressing the formation of stress granules. Signal Transduct Target Ther. 2022;7(1):22.

51. Nakagawa K, Narayanan K, Wada M, Makino S. Inhibition of Stress Granule Formation by Middle East Respiratory Syndrome Coronavirus 4a Accessory Protein Facilitates Viral Translation, Leading to Efficient Virus Replication. J Virol. 2018;92(20).

52. Khandjian EW, Bardoni B, Corbin F, Sittler A, Giroux S, Heitz D, et al. Novel Isoforms of the Fragile X Related Protein FXR1P are Expressed During Myogenesis. Hum Mol Genet. 1998;7(13):2121–8.

53. Ho W, Huang J. The return of the rings: Evolutionary convergence of aromatic residues in the intrinsically disordered regions of RNA-binding proteins for liquid–liquid phase separation. Protein Sci. 2022;31(5):e4317.

54. Lin Y, Currie SL, Rosen MK. Intrinsically disordered sequences enable modulation of protein phase separation through distributed tyrosine motifs. J Biol Chem. 2017;292(46):19110–20.

55. Meshram CD, Phillips AT, Lukash T, Shiliaev N, Frolova EI, Frolov I. Mutations in Hypervariable Domain of Venezuelan Equine Encephalitis Virus nsP3 Protein Differentially Affect Viral Replication. J Virol. 2020;94(3).

56. Meshram CD, Shiliaev N, Frolova EI, Frolov I. Hypervariable Domain of nsP3 of Eastern Equine Encephalitis Virus Is a Critical Determinant of Viral Virulence. J Virol. 2020;94(17).

57. Oh SW, Onomoto K, Wakimoto M, Onoguchi K, Ishidate F, Fujiwara T, et al. Leader-Containing Uncapped Viral Transcript Activates RIG-I in Antiviral Stress Granules. PLoS Pathog. 2016;12(2):e1005444.

58. Paget M, Cadena C, Ahmad S, Wang HT, Jordan TX, Kim E, et al. Stress granules are shock absorbers that prevent excessive innate immune responses to dsRNA. Mol Cell. 2023;83(7):1180–1196.e8.

59. Luo L, Li Z, Zhao T, Ju X, Ma P, Jin B, et al. SARS-CoV-2 nucleocapsid protein phase separates with G3BPs to disassemble stress granules and facilitate viral production. Sci Bull. 2021;66(12):1194–204.

60. Nabeel-Shah S, Lee H, Ahmed N, Burke GL, Farhangmehr S, Ashraf K, et al. SARS-CoV-2 nucleocapsid protein binds host mRNAs and attenuates stress granules to impair host stress response. Iscience. 2022;25(1):103562.

61. Zheng ZQ, Wang SY, Xu ZS, Fu YZ, Wang YY. SARS-CoV-2 nucleocapsid protein impairs stress granule formation to promote viral replication. Cell Discov. 2021;7(1):38.

62. Luo L, Li Z, Ma P, Zou Y, Li P, Liang A, et al. SARS-CoV-2 Nucleocapsid Protein Impairs SG Assembly by Partitioning into G3BP Condensate. SSRN Electron J. 2020;

63. Zhao Y, Sui L, Wu P, Wang W, Wang Z, Yu Y, et al. A dual-role of SARS-CoV-2 nucleocapsid protein in regulating innate immune response. Signal Transduct Target Ther. 2021;6(1):331.

64. Cong Y, Ulasli M, Schepers H, Mauthe M, V’kovski P, Kriegenburg F, et al. Nucleocapsid Protein Recruitment to Replication-Transcription Complexes Plays a Crucial Role in Coronaviral Life Cycle. J Virol. 2020;94(4):e01925–19.

65. Carlson CR, Asfaha JB, Ghent CM, Howard CJ, Hartooni N, Safari M, et al. Phosphoregulation of Phase Separation by the SARS-CoV-2 N Protein Suggests a Biophysical Basis for its Dual Functions. Mol Cell. 2020;80(6):1092–1103.e4.

66. Bai Z, Cao Y, Liu W, Li J. The SARS-CoV-2 Nucleocapsid Protein and Its Role in Viral Structure, Biological Functions, and a Potential Target for Drug or Vaccine Mitigation. Viruses. 2021;13(6):1115.

67. Lu S, Ye Q, Singh D, Cao Y, Diedrich JK, Yates JR, et al. The SARS-CoV-2 nucleocapsid phosphoprotein forms mutually exclusive condensates with RNA and the membrane-associated M protein. Nat Commun. 2021;12(1):502.

68. Scherer KM, Mascheroni L, Carnell GW, Wunderlich LCS, Makarchuk S, Brockhoff M, et al. SARS-CoV-2 nucleocapsid protein adheres to replication organelles before viral assembly at the Golgi/ERGIC and lysosome-mediated egress. Sci Adv. 2022;8(1):eabl4895.

69. Harcourt J, Tamin A, Lu X, Kamili S, Sakthivel SK, Murray J, et al. Severe Acute Respiratory Syndrome Coronavirus 2 from Patient with Coronavirus Disease, United States - Volume 26, Number 6—June 2020 - Emerging Infectious Diseases journal - CDC. Emerg Infect Dis. 2020;26(6):1266–73.

70. Holshue ML, DeBolt C, Lindquist S, Lofy KH, Wiesman J, Bruce H, et al. First Case of 2019 Novel Coronavirus in the United States. N Engl J Med. 2020;382(10):929–36.

71. Johnson BA, Xie X, Bailey AL, Kalveram B, Lokugamage KG, Muruato A, et al. Loss of furin cleavage site attenuates SARS-CoV-2 pathogenesis. Nature. 2021;591(7849):293–9.

72. Johnson BA, Zhou Y, Lokugamage KG, Vu MN, Bopp N, Crocquet-Valdes PA, et al. Nucleocapsid mutations in SARS-CoV-2 augment replication and pathogenesis. Plos Pathog. 2022;18(6):e1010627.

73. Lamichhane PP, Aditi, Xie X, Samir P. Cell-Type-Specific Effect of Innate Immune Signaling on Stress Granules. Stresses. 2024;4(3):411–20.

74. Mazo G. QuickFigures: A toolkit and ImageJ PlugIn to quickly transform microscope images into scientific figures. PLoS ONE. 2021;16(11):e0240280.

